# Genetic diversity and population structure of Ethiopian taro (*Colocasia esculenta* (L.) Schott) as revealed by SSR markers

**DOI:** 10.64898/2026.10.28.762411

**Authors:** Shimekit Tadele, Tilahun Mekonnen, Kassahun Tesfaye, Sebsebe Demissew, Wendawek Abebe, Ermias Lulekal, Mulugeta Kebede, Zelalem Gebremariam, Asfaw Kifle, Agegnehu Wasse, Alok Kumar, Abdi Mohammed, Mulatu Geleta

## Abstract

Taro (*Colocasia esculenta* (L.) Schott) is an important root and tuber crop that contributes to food security and livelihoods, but its genetic diversity remains poorly characterized in Ethiopia. This study assessed Ethiopian taro genetic diversity and population structure using simple sequence repeat (SSR) markers. We analyzed 119 accessions, 79 collected from farmers’ fields and 40 from the Ethiopian Biodiversity Institute (EBI), representing three regional populations: South Ethiopia Regional State (SERS), South West Ethiopia Peoples’ Regional State (SWEPRS), and Oromia. Fifteen polymorphic SSR markers produced 199 reproducible polymorphic loci, with 100% polymorphism. Polymorphism information content (PIC) ranged from 0.300 to 0.888 (mean 0.583). Diversity based on scored binary marker states was relatively low, with a mean effective number of alleles (Ne) of 1.077 and expected heterozygosity (He) of 0.066. Across populations, He ranged from 0.047 to 0.061, and the percentage of polymorphic loci ranged from 37.14% to 82.57%. Analysis of molecular variance showed that 98% of molecular variation occurred within populations and 2% among populations. Genetic differentiation (ΦPT, proportion of molecular variation among populations) was weak but significant (ΦPT = 0.016, P = 0.016). Principal coordinate analysis, neighbor-joining (NJ), and unweighted pair-group method with arithmetic mean (UPGMA) clustering showed extensive overlap among accessions and no clear geographic structuring. Bayesian clustering identified two groups as the strongest partition according to the ΔK criterion, but substantial admixture occurred, and groups did not correspond clearly to geographic populations. Despite relatively low diversity at individual scored loci, all 119 accessions had unique multilocus genotypes, indicating high multilocus genotypic diversity. Most molecular variation occurred within rather than among regional populations, potentially reflecting planting-material exchange and predominantly clonal propagation. These findings provide a basis for conserving, characterizing, and utilizing Ethiopian taro germplasm in crop improvement.

## Introduction

Root and tuber crops play a crucial role in ensuring food security and generating income for rural households across Africa [1]. Among these, taro (*Colocasia esculenta* (L.) Schott), a member of the family Araceae and sub-family Aroideae, is one of the oldest cultivated food crops known to humankind [2–5]. Archaeological evidence suggests that its utilization dates back approximately 28,000 years, with at least 2,000 years of domestication history [4,6]. Taro is a vegetatively propagated, perennial monocotyledonous plant characterized by underground corms, although it is typically cultivated as an annual crop [7] and is well adapted to diverse agro-ecological conditions [2,8,9].

Historically, before the Columbian exchange, *C. esculenta* was among the most widely distributed food crops, cultivated across tropical and temperate regions including Africa, Asia, the Mediterranean, and Oceania. Its origin is generally associated with tropical Asia, with subsequent dissemination to Africa possibly through the Mediterranean region, reaching Egypt around 2000 BC and spreading further via trade and migration routes [10]. Today, taro is cultivated globally, with major production areas in Africa, Southeast Asia, and the Pacific, particularly in countries such as China, Japan, and several island nations [11]. In Africa, it holds significant economic and nutritional value, especially in countries like Nigeria, Cameroon, Ghana, and Burkina Faso [12], and contributes to both subsistence and commercial agriculture in Ethiopia..

Taro is consumed by billions of people worldwide [13] and can be processed into a wide range of food products, including boiled, roasted, or fried corms, flour, fermented products, and chips [14]. Its leaves are also nutritionally valuable, being rich in vitamins, minerals, fiber, and bioactive compounds [14]. Beyond its nutritional role, taro has medicinal applications in traditional practices, where it is used to treat conditions such as ulcers, tuberculosis, and fungal infections in several African countries, including Ethiopia [15]. Despite these benefits, taro remains categorized as an underutilized or orphan crop [13], receiving comparatively limited research attention relative to other staple crops.

Globally, taro ranks 14^th^ among staple crops and is the fifth most consumed root crop [15,16]. However, its genetic improvement has been constrained by limited research efforts, insufficient characterization of germplasm, and a lack of well-developed breeding programs [17,18]. Furthermore, taro genetic resources are increasingly threatened by genetic erosion due to changes in cropping systems, replacement by improved varieties, and shifts in agricultural practices [1]. In Ethiopia, analysis of long-term production data indicates that increases in taro output are primarily driven by expansion of cultivated area rather than improvements in yield, highlighting the need for genetic enhancement strategies.

Cytogenetically, *C. esculenta* is regarded as a single polymorphic species exhibiting variation in ploidy levels [13]. Both diploid (2n = 2x = 28) and triploid (2n = 3x = 42) forms are reported, with diploids predominating in certain regions and triploids more common in others, particularly at higher altitudes and latitudes in Asia [10,19]. Two major taxonomic groups are recognized: the dasheen type (*C. esculenta* var. esculenta), characterized by a large central corm, and the eddoe type (*C. esculenta* var. antiquorum), which produces numerous smaller cormels [20]. This cytological and morphological diversity underscores the importance of detailed genetic characterization for effective utilization and conservation.

Molecular marker technologies have become essential tools for assessing genetic diversity, population structure, and phylogenetic relationships in crop species [21,22]. Earlier studies on taro employed techniques such as isozymes [23], amplified fragment length polymorphism (AFLP) [24], and random amplified polymorphic DNA (RAPD) [25]. While these methods provided valuable insights, they are often limited by issues of reproducibility and resolution. In contrast, Simple sequence repeat (SSR) markers have emerged as highly reliable and informative tools due to their codominant inheritance, high polymorphism, reproducibility, and genome-wide distribution [3,26–28].

SSRs have been successfully applied in taro for cultivar identification, assessment of genetic diversity, and analysis of population structure [15,29,30]. These markers enable precise detection of allelic variation and facilitate the selection of informative loci across the genome, making them particularly suitable for germplasm characterization and breeding applications [31]. Moreover, SSR-based studies have demonstrated their effectiveness in distinguishing taro genotypes and identifying genetic relationships among accessions [21,32]. Although more recent marker systems such as single-nucleotide polymorphisms (SNPs) and DArTseq have been introduced [33], SSRs remain widely preferred due to their cost-effectiveness and analytical robustness.

Despite these advancements, comprehensive studies on taro genetic diversity, particularly in underrepresented regions, remain limited. A thorough understanding of genetic variability and population structure is essential for germplasm conservation, breeding program development, and the identification of cultivars with desirable traits such as stress tolerance and disease resistance [34,35]. Therefore, the present study aims to assess the genetic diversity and population structure of Ethiopian taro cultivars using SSR markers, with the objective of generating valuable information for effective conservation and genetic improvement of this important yet underutilized crop.

## Materials and methods

### Plant materials

A total of 119 accessions used in this study, of which 79 accessions collected from farmers’ fields while 40 accessions obtained from Ethiopian Biodiversity Institute (EBI). Overall, the collection sites covered three regional states of Ethiopia, which were treated as three populations in the analysis: South West Ethiopia Peoples’ Regional State (SWEPRS), comprising Kaffa, Dawuro, Bench Maji, and Sheka zones; South Ethiopia Regional State (SERS), comprising Wolaita and Gedeo zones; and Oromiya, comprising Jimma, Illubabor, East Wellega, and West Wellega zones (Fig 1). Full passport data for the accessions are provided in S1 Table. The collections were planted in pots in greenhouse of Institute of Biotechnology at Arat Kilo Campus of Addis Ababa University (AAU). Clean young leaves were collected and dried using silica gel and kept at room temperature for DNA extraction.

**Fig 1.** Map of the study area illustrating the sampling sites included in the study.

### Genomic DNA extraction

Genomic DNA was extracted from dried leaf samples (0.4 g per sample) using the Cetyltrimethylammonium bromide (CTAB) protocol optimized by [36] at the Plant Genetics Research Laboratory, Addis Ababa University (AAU). The extracted DNA was stored at −20°C until polymerase chain reaction (PCR) analysis. DNA quality and integrity were assessed by electrophoresis on a 1% agarose gel stained with ethidium bromide. DNA concentration and purity were evaluated using a NanoDrop ND-2000 spectrophotometer (Thermo Fisher Scientific, Wilmington, DE, USA).

### Selection of molecular markers and PCR amplification

Based on the strength of informativeness, Taro SSR markers previously used by the researchers were selected for in this study [1,6,31,37–40]. A total of 25 SSR markers were used in preliminary screening, of which 15 that produced clear, consistent, and reproducible amplification profiles were selected for genotyping (Table 1). PCR amplification was conducted at the Biotechnology Laboratory of the School of Plant Science, Haramaya University, Ethiopia.

**Table 1.**
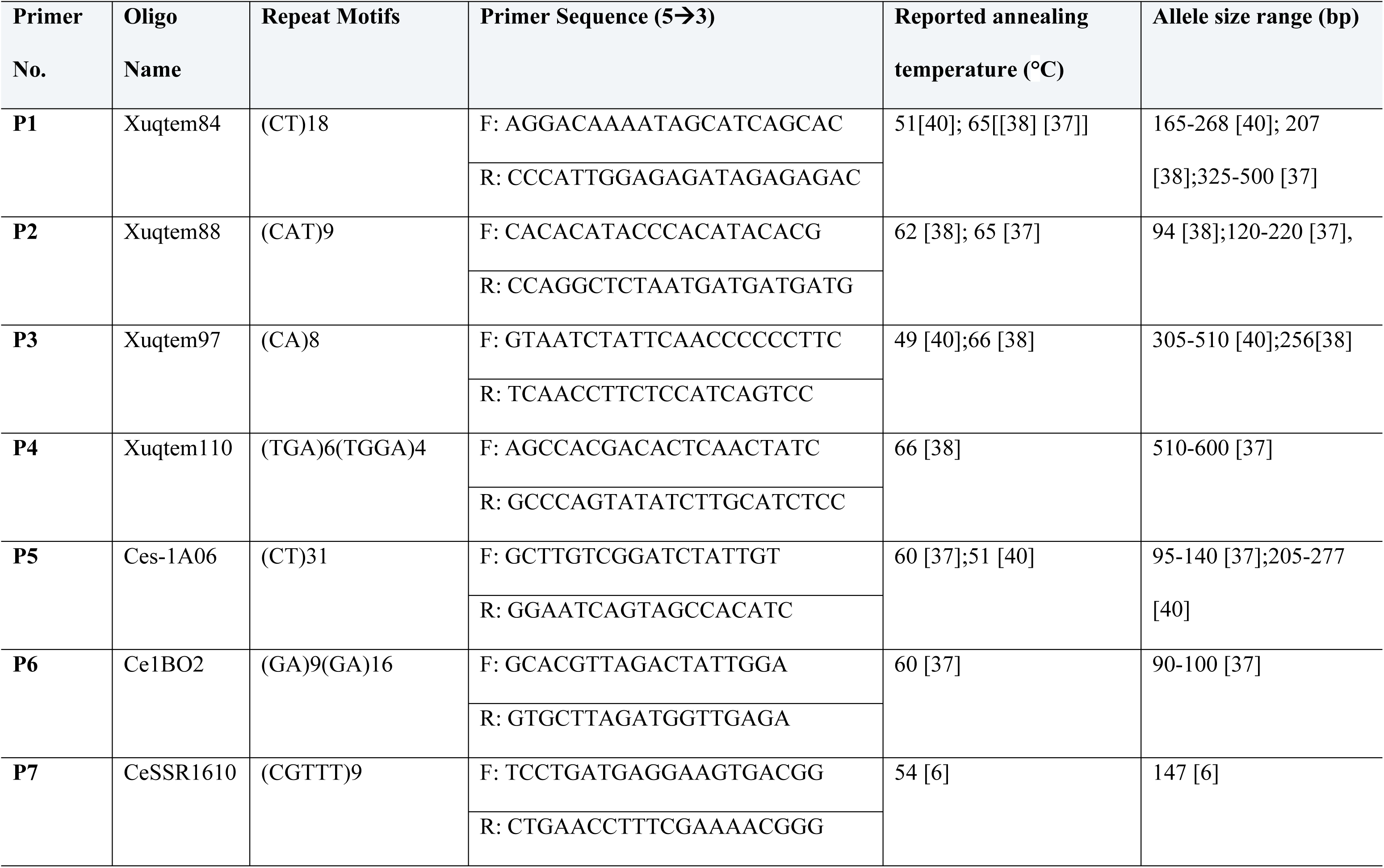
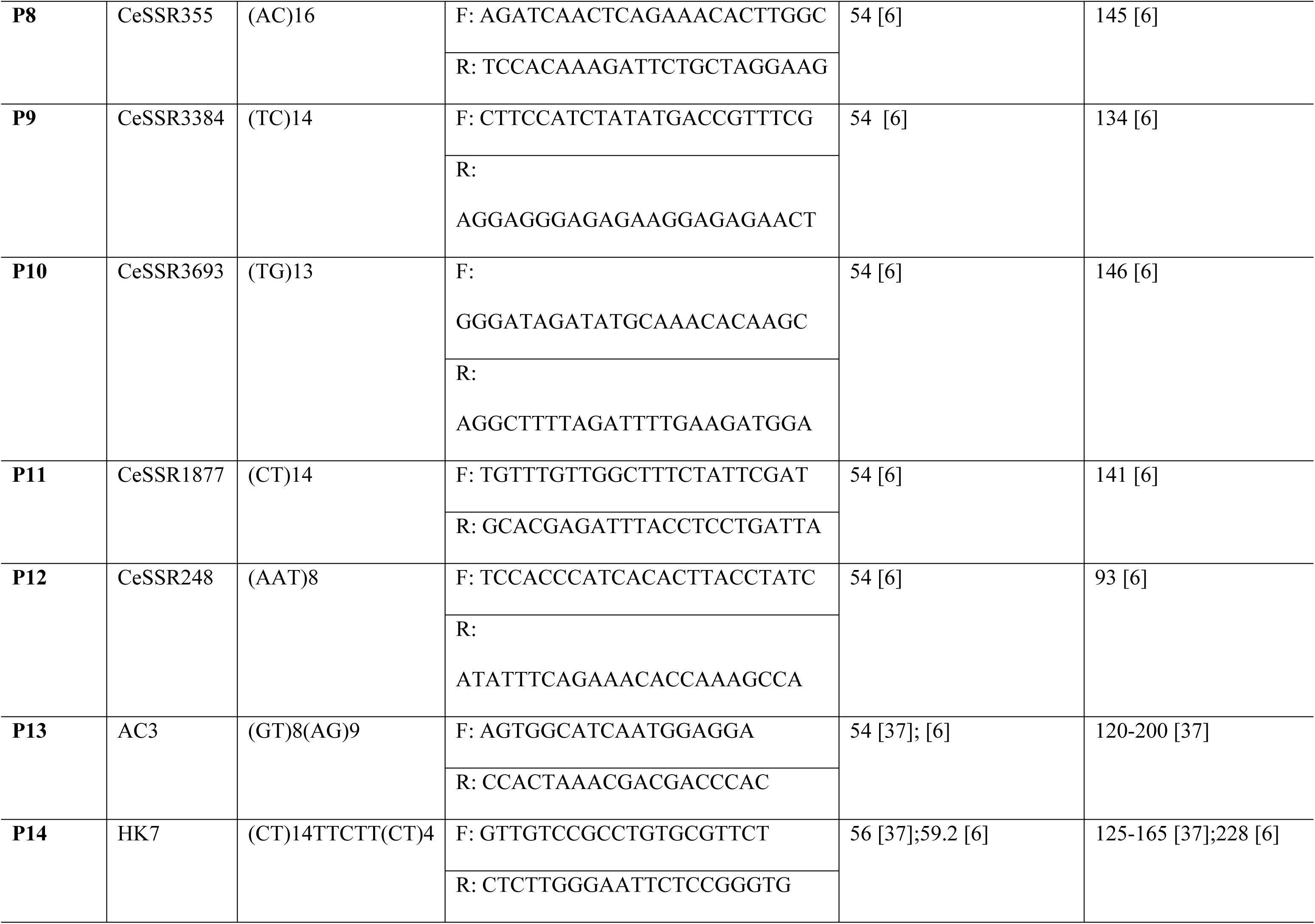
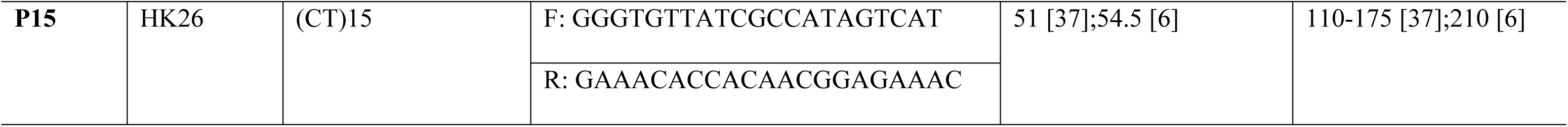
Characteristics of the SSR markers used for genetic analysis of Ethiopian taro (*C. esculenta*) germplasm.

PCR reactions were performed using an Eppendorf Flexlid thermal cycler (96 wells) in a final reaction volume of 20 μL containing 2 μL 10x reaction buffer, 1.2 μL MgCl₂, 0.6 μL of each primer, 0.1 μL Taq polymerase, and 2 μL template DNA (20 ng/μL) with the final volume adjusted to 20 μL. The PCR profile consisted of an initial denaturation at 95°C for 2 min, followed by 35 cycles of denaturation at 95°C for 1 min, primer-specific annealing temperatures ranging from 45–53°C for 1 min (Table 1), and extension at 72°C for 2 min, with a final extension at 72°C for 10 min and holding at 10°C.

Amplified products were separated on 2% agarose gel stained with ethidium bromide (10 mg/mL) prepared in 0.5x Tris-borate-EDTA buffer (TBE) buffer. Electrophoresis was conducted using a horizontal mini-gel electrophoresis unit (Cole-Parmer™, Fisher Scientific, Catalog No. 28560-10) at 120 V for 1 h 20 min. A 100 bp DNA ladder (BioBase) was used as a molecular size standard. PCR products were visualized and photographed using a UVP BioDoc-It™ imaging system (USA).

### Data analysis

Band size estimation and migration distance measurements were performed using Fiji Is Just ImageJ software (ImageJ version 2.16.0). Clear and reproducible SSR loci were scored using a ±3 bp binning approach. Strong and distinct bands were scored as present (1) or absent (0), while ambiguous bands were excluded. Because *C. esculenta* exhibits complex ploidy, accurate determination of allele dosage from SSR banding patterns is difficult. Therefore, SSR profiles were converted into binary matrix, with each scored band treated as an independent dominant locus for subsequent analyses [4,41,42]. The resulting binary matrix was used for downstream genetic analyses.

The binary dataset was analyzed using GenAlEx version 6.5 [43] with the dominant-marker option. Genetic diversity parameters, including the number of alleles (Na), effective number of alleles (Ne), Shannon’s information index (I), expected heterozygosity (He), and unbiased expected heterozygosity (uHe), were estimated under Hardy–Weinberg equilibrium assumptions. Analysis of molecular variance (AMOVA), pairwise genetic differentiation (ΦPT), and principal coordinates analysis (PCoA) were performed in GenAlEx to assess the partitioning of genetic variation and relationships among accessions and regional populations.

Additional genetic diversity and multilocus analyses were performed in R version 4.5.2 using the adegenet and poppr packages. Genotypic diversity was assessed using the number of multilocus genotypes (MLGs), expected multilocus genotype richness (eMLG), Shannon–Wiener diversity index (H), Stoddart and Taylor’s index of genotypic diversity (G), Simpson’s index of diversity (λ), genotypic evenness (E.5), and expected heterozygosity (Hexp). Multilocus association was evaluated using the index of association (Ia) and the standardized index of association (r̄d). Genetic relationships among accessions were further assessed using Nei’s genetic distances and Neighbor-Joining (NJ) and unweighted pair-group method with arithmetic mean (UPGMA) clustering methods implemented in R.

Prior to downstream analyses, markers and samples with more than 20% missing data were excluded to improve data quality and analytical reliability. Consequently, the initial dataset of 182 taro accessions genotyped with 25 SSR markers was filtered to retain 119 accessions and 15 high-quality, reproducible SSR markers for subsequent analyses.

Population structure was assessed using STRUCTURE version 2.3.4 under an admixture model with correlated allele frequencies. The analysis was performed for K = 1–10, with five independent runs conducted for each K value. Each run consisted of 100,000 iterations following a burn-in period of 50,000 iterations. The most strongly supported number of genetic clusters was evaluated using the Evanno ΔK method [44], based on the rate of change in the log probability of the data between successive K values. The STRUCTURE results were visualized as individual membership coefficients (Q-values), with accessions displayed according to their predefined regional populations.

## Results

### SSR amplification profile

The A260/A280 ratios of the extracted DNA samples ranged from 1.78 to 2.07, while the A260/A230 ratios ranged from 1.98 to 2.16. These measurements indicated that the extracted DNA was of adequate purity for subsequent SSR analysis.

Fifteen SSR primers generated clear and reproducible amplification profiles across the taro germplasm collections. The amplified DNA fragments ranged from 60 bp to 490 bp, which was broader than the previously reported range of 90–287 bp [45]. Following scoring and data quality filtering, a final dataset consisting of 119 accessions, 15 SSR markers, and three population groups was retained for downstream analyses.

### Genetic informativeness and polymorphism of SSR markers

The genetic informativeness and polymorphism parameters of the 15 SSR primers are presented in Table 2. Because the SSR profiles were analyzed as dominant presence/absence data, the diversity statistics estimated by GenAlEx (Na, Ne, I, He, and uHe) represent variation in the binary marker dataset, with Na reflecting the mean number of observed allelic states per scored locus rather than allele counts from codominant SSR genotypes. Across the 15 primers, a total of 199 loci were amplified, and all loci were polymorphic, resulting in 100% polymorphism for each primer.

**Table 2.** Genetic informativeness and polymorphism of SSR primers.

| Primer | Annealing temperature (°C) | Total Loci | Polymorphic Loci | % Polymorphism | Allele size range (bp) | Na (Mean ± SE) | Ne (Mean ± SE) | I (Mean ± SE) | He (Mean ± SE) | uHe (Mean ± SE) | PIC |
|---|---|---|---|---|---|---|---|---|---|---|---|
| **P1** | 45 | 15 | 15 | 100 | 135–225 | 1.911 ± 0.008 | 1.184 ± 0.005 | 0.270 ± 0.005 | 0.148 ± 0.003 | 0.151 ± 0.003 | 0.888 ± 0.0012 |
| **P2** | 45 | 15 | 15 | 100 | 200–490 | 1.288 ± 0.053 | 1.056 ± 0.003 | 0.105 ± 0.005 | 0.050 ± 0.003 | 0.051 ± 0.003 | 0.300 ± 0.0012 |
| **P3** | 46 | 14 | 14 | 100 | 150–465 | 1.211 ± 0.047 | 1.042 ± 0.002 | 0.082 ± 0.004 | 0.038 ± 0.002 | 0.038 ± 0.002 | 0.492 ± 0.0008 |
| **P4** | 46 | 13 | 13 | 100 | 190–300 | 1.846 ± 0.014 | 1.081 ± 0.003 | 0.153 ± 0.004 | 0.072 ± 0.002 | 0.074 ± 0.002 | 0.444 ± 0.0008 |
| **P5** | 46 | 14 | 14 | 100 | 200–350 | 1.248 ± 0.052 | 1.036 ± 0.002 | 0.077 ± 0.004 | 0.034 ± 0.002 | 0.034 ± 0.002 | 0.462 ± 0.0008 |
| **P6** | 50 | 14 | 14 | 100 | 185–260 | 1.429 ± 0.044 | 1.055 ± 0.003 | 0.108 ± 0.005 | 0.050 ± 0.002 | 0.052 ± 0.002 | 0.648 ± 0.0008 |
| **P7** | 47 | 13 | 13 | 100 | 85–170 | 1.179 ± 0.062 | 1.069 ± 0.004 | 0.122 ± 0.007 | 0.061 ± 0.003 | 0.063 ± 0.004 | 0.732 ± 0.0012 |
| **P8** | 46 | 14 | 14 | 100 | 120–160 | 1.238 ± 0.062 | 1.062 ± 0.004 | 0.109 ± 0.006 | 0.054 ± 0.003 | 0.055 ± 0.003 | 0.540 ± 0.0012 |
| **P9** | 46 | 17 | 17 | 100 | 150–400 | 0.921 ± 0.036 | 1.036 ± 0.002 | 0.069 ± 0.003 | 0.032 ± 0.002 | 0.033 ± 0.002 | 0.414 ± 0.0008 |
| **P10** | 53 | 14 | 14 | 100 | 130–200 | 1.381 ± 0.040 | 1.051 ± 0.002 | 0.102 ± 0.004 | 0.046 ± 0.002 | 0.047 ± 0.002 | 0.612 ± 0.0008 |
| **P11** | 56 | 11 | 11 | 100 | 110–150 | 1.576 ± 0.039 | 1.116 ± 0.006 | 0.169 ± 0.007 | 0.090 ± 0.004 | 0.092 ± 0.004 | 0.540 ± 0.0016 |
| **P12** | 46 | 12 | 12 | 100 | 60–110 | 1.500 ± 0.031 | 1.086 ± 0.004 | 0.147 ± 0.006 | 0.074 ± 0.003 | 0.075 ± 0.003 | 0.444 ± 0.0012 |
| **P13** | 50 | 16 | 16 | 100 | 120–400 | 1.475 ± 0.038 | 1.071 ± 0.003 | 0.130 ± 0.005 | 0.063 ± 0.002 | 0.064 ± 0.002 | 0.780 ± 0.0008 |
| **P14** | 47 | 8 | 8 | 100 | 140–160 | 1.733 ± 0.025 | 1.137 ± 0.004 | 0.216 ± 0.005 | 0.114 ± 0.003 | 0.116 ± 0.003 | 0.696 ± 0.0012 |
| **P15** | 53 | 9 | 9 | 100 | 150–180 | 1.333 ± 0.046 | 1.080 ± 0.003 | 0.126 ± 0.005 | 0.065 ± 0.002 | 0.066 ± 0.003 | 0.756 ± 0.0008 |
| **Mean** | — | 13.27 | 13.27 | 100 | — | 1.418 ± 0.069 | 1.077 ± 0.011 | 0.132 ± 0.014 | 0.066 ± 0.008 | 0.067 ± 0.008 | 0.583 ± 0.001 |

**Note:** Na = number of observed allelic states; Ne = effective number of alleles; I = Shannon's information index; He = expected heterozygosity; uHe = unbiased expected heterozygosity; SE = standard error.

The mean number of observed allelic states per locus (Na) was 1.418 ± 0.069, while the effective number of alleles (Ne) averaged 1.077 ± 0.011. Shannon’s information index (I) ranged from 0.069 ± 0.003 to 0.270 ± 0.005, with an overall mean of 0.132 ± 0.014. Expected heterozygosity (He) ranged from 0.032 ± 0.002 to 0.148 ± 0.003, averaging 0.066 ± 0.008, whereas unbiased expected heterozygosity (uHe) ranged from 0.033 ± 0.002 to 0.151 ± 0.003, with a mean of 0.067 ± 0.008. Polymorphism information content (PIC) varied among primers, ranging from 0.300 ± 0.0012 to 0.888 ± 0.0012, with an overall mean of 0.583 ± 0.001. The allele-size ranges varied among primers, from 60–110 bp for P12 to 200–490 bp for P2. Overall, all 15 primers produced polymorphic loci across the analyzed accessions, although the diversity estimates varied among primers.

### Population genetic diversity

The genetic diversity parameters of the three regional populations are presented in Table 3. The populations comprised 44 accessions from SERS, 59 from SWEPRS, and 16 from Oromia. The mean number of observed allelic states (Na) ranged from 0.743 ± 0.052 in Oromia to 1.651 ± 0.041 in SWEPRS, whereas the effective number of alleles (Ne) showed relatively little variation, ranging from 1.057 ± 0.005 in Oromia to 1.069 ± 0.004 in SERS.

**Table 3.** Genetic diversity parameters of the three administrative-region populations.

| **Population** | **N** | **Na** | **Ne** | **Shannon's I** | **He** | **uHe** | **Polymorphic loci (%)** |
|---|---|---|---|---|---|---|---|
| **SERS** | 44 | 1.531 ± 0.045 | 1.069 ± 0.004 | 0.126 ± 0.006 | 0.061 ± 0.003 | 0.061 ± 0.003 | 76.57 |
| **SWEPRS** | 59 | 1.651 ± 0.041 | 1.061 ± 0.003 | 0.119 ± 0.005 | 0.055 ± 0.003 | 0.056 ± 0.003 | 82.57 |
| **Oromia** | 16 | 0.743 ± 0.052 | 1.057 ± 0.005 | 0.089 ± 0.007 | 0.047 ± 0.004 | 0.048 ± 0.004 | 37.14 |
| **Overall mean ± SE** | — | 1.309 ± 0.029 | 1.062 ± 0.002 | 0.111 ± 0.003 | 0.054 ± 0.002 | 0.055 ± 0.002 | 65.43 ± 14.25 |

Note: N represents the actual number of accessions sampled in each administrative-region population. Na, Ne, I, He, and uHe are locus-averaged diversity estimates.

Shannon’s information index (I) ranged from 0.089 ± 0.007 in Oromia to 0.126 ± 0.006 in SERS, while expected heterozygosity (He) ranged from 0.047 ± 0.004 in Oromia to 0.061 ± 0.003 in SERS. Unbiased expected heterozygosity (uHe) ranged from 0.048 ± 0.004 in Oromia to 0.061 ± 0.003 in SERS. SWEPRS showed the highest Na (1.651 ± 0.041) and the highest proportion of polymorphic loci (82.57%), whereas Oromia showed the lowest values for Na, I, He, uHe, and the proportion of polymorphic loci. SERS exhibited the highest I and He values. Across the three populations, the proportion of polymorphic loci ranged from 37.14% in Oromia to 82.57% in SWEPRS, with an overall mean of 65.43 ± 14.25%.

### Population-level band profiles

The population-level distribution of amplified bands and private bands is presented in Table 4. SWEPRS had the highest total number of bands (289), followed by SERS (268), whereas Oromia had substantially fewer bands (130). SWEPRS also possessed the highest number of private bands (44), followed by SERS (30), while Oromia had three private bands. The number of bands occurring at frequencies of ≥5% was 176 in SERS, 174 in SWEPRS, and 130 in Oromia. The corresponding mean He values were 0.061 ± 0.003, 0.055 ± 0.003, and 0.047 ± 0.004, respectively.

**Table 4.** Population-level polymorphism and band characteristics.

| **Population** | **Total bands** | **Bands ≥5% frequency** | **Private bands** | **Mean He ± SE** | **Mean uHe ± SE** |
|---|---|---|---|---|---|
| **SERS** | 268 | 176 | 30 | 0.061 ± 0.003 | 0.061 ± 0.003 |
| **SWEPRS** | 289 | 174 | 44 | 0.055 ± 0.003 | 0.056 ± 0.003 |
| **Oromia** | 130 | 130 | 3 | 0.047 ± 0.004 | 0.048 ± 0.004 |

The Binary Allelic Pattern (BAPT) analysis indicated that no locally common bands were detected at the ≤25% or ≤50% frequency categories.

### Genetic distance and genetic identity among populations

Pairwise Nei’s genetic distance and genetic identity among the three regional populations are presented in Table 5. Nei’s genetic distances were very low, ranging from 0.001 to 0.002, while genetic identity values ranged from 0.998 to 0.999. The smallest genetic distance was observed between SERS and SWEPRS (D = 0.001; I = 0.999), whereas SERS–Oromia and SWEPRS–Oromia each showed a distance of 0.002 and genetic identity of 0.998.

**Table 5.** Pairwise Nei’s genetic distance and genetic identity among administrative-region populations.

| **Population pair** | **Nei's genetic distance (D)** | **Nei's genetic identity (I)** |
|---|---|---|
| **SERS – SWEPRS** | 0.001 | 0.999 |
| **SERS – Oromia** | 0.002 | 0.998 |
| **SWEPRS – Oromia** | 0.002 | 0.998 |

UPGMA clustering based on the mean pairwise genetic distances among the three regional populations is presented in Fig 2. SERS and SWEPRS formed the closest cluster, whereas Oromia was comparatively more distant from the other two populations.

**Fig 2.**
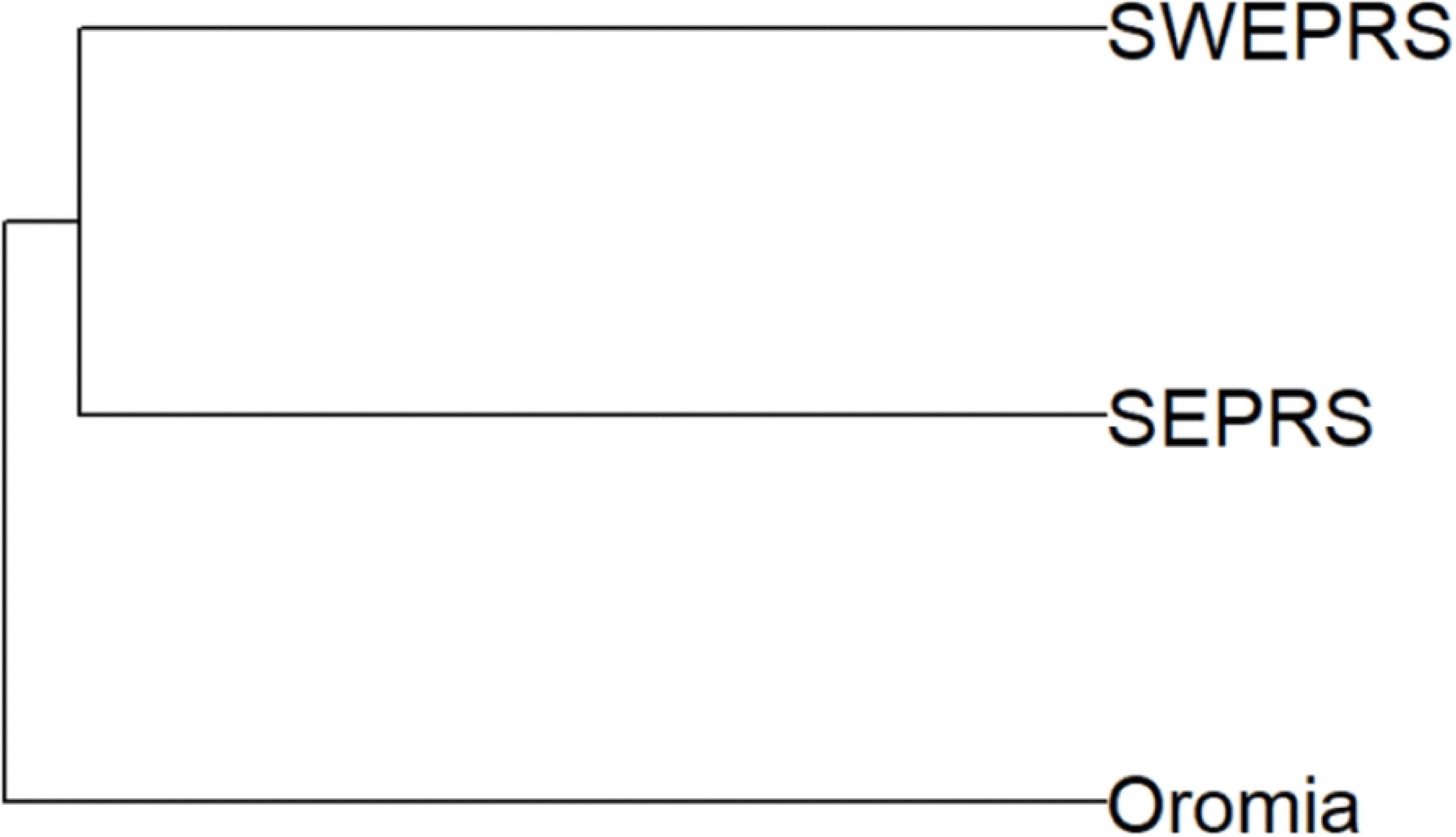
UPGMA phylogenetic clustering of the three regional populations (SWEPRS, SERS and Oromia) based on mean pairwise genetic distance. *Note: SEPRS in the figure refers to SERS*.

### Analysis of molecular variance

AMOVA partitioned 98% of the total molecular variation within populations and 2% among populations (Table 6; Fig 3). The estimated variance components were 47.998 within populations and 0.775 among populations, with a total estimated variance of 48.772. The overall ΦPT was 0.016 and was statistically significant (*P* = 0.016), indicating weak but detectable differentiation among the three regional populations.

**Fig 3.**
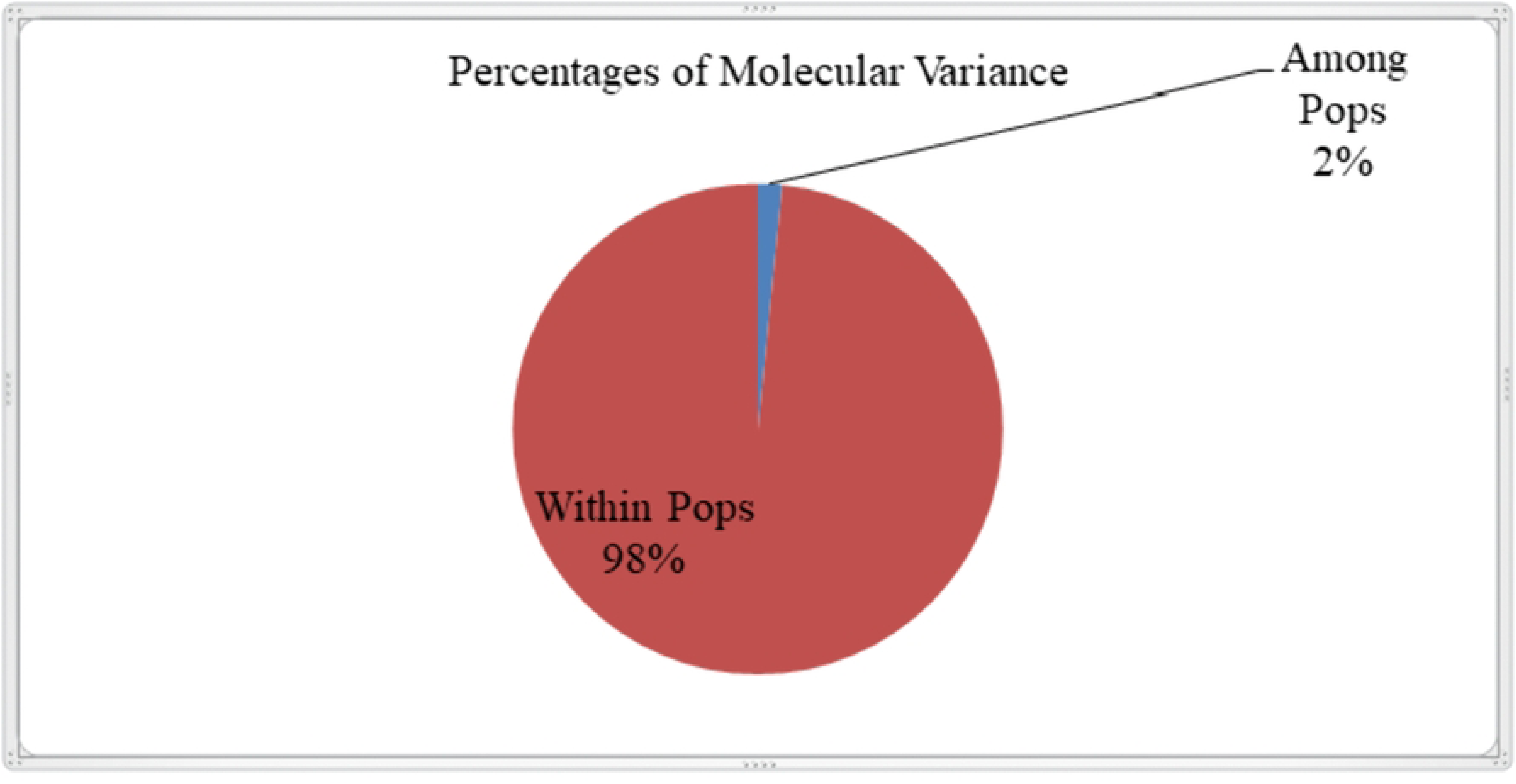
AMOVA showing the partitioning of genetic variation among and within taro (*C. esculenta*) populations based on SSR marker data.

**Table 6.** AMOVA among and within the three regional populations of taro.

| **Source of variation** | **df** | **SS** | **MS** | **Estimated variance** | **Variation (%)** |
|---|---|---|---|---|---|
| **Among populations** | 2 | 151.244 | 75.622 | 0.775 | 2 |
| **Within populations** | 116 | 5567.714 | 47.998 | 47.998 | 98 |
| **Total** | 118 | 5718.958 | — | 48.772 | 100 |

Overall ΦPT = 0.016; P = 0.016.

### Pairwise genetic differentiation and gene flow

Pairwise genetic differentiation and estimated gene flow among the three regional populations are presented in Table 7. ΦPT values were low, ranging from 0.010 between SERS and SWEPRS to 0.038 between SERS and Oromia. Significant differentiation was detected only between SERS and Oromia (ΦPT = 0.038, *P* = 0.018). The differentiation between SERS and SWEPRS (ΦPT = 0.010, *P* = 0.074) and between SWEPRS and Oromia (ΦPT = 0.015, *P* = 0.127) was not statistically significant.

**Table 7.** Pairwise genetic differentiation (ΦPT), significance, linearized ΦPT, and Nm among the three regional populations of taro.

| **Population pair** | **ΦPT** | **P-value** | **Significance** | **Linearized ΦPT** | **Nm** |
|---|---|---|---|---|---|
| **SERS–SWEPRS** | 0.010 | 0.074 | NS | 0.010 | 24.750 |
| **SERS–Oromia** | 0.038 | 0.018 | * | 0.040 | 6.342 |
| **SWEPRS–Oromia** | 0.015 | 0.127 | NS | 0.015 | 16.417 |

NS = not significant; *P < 0.05. ΦPT = proportion of molecular variation attributable to differences among populations; Nm = estimated gene flow, calculated as Nm = (1 − ΦPT)/(4ΦPT).

Estimated gene flow was high among all population pairs, with Estimated gene-flow (Nm) values ranging from 6.342 between SERS and Oromia to 24.750 between SERS and SWEPRS. The estimated Nm between SWEPRS and Oromia was 16.417.

### Principal coordinates analysis

The PCoA based on the genetic distance matrix showed that the first, second, and third coordinates explained 9.30%, 5.27%, and 4.76% of the total variation, respectively, with the first three axes cumulatively accounting for 19.32% of the variation (Fig 4). The accessions showed extensive overlap among SERS, SWEPRS, and Oromia, with no clear separation according to geographic origin.

**Fig 4.**
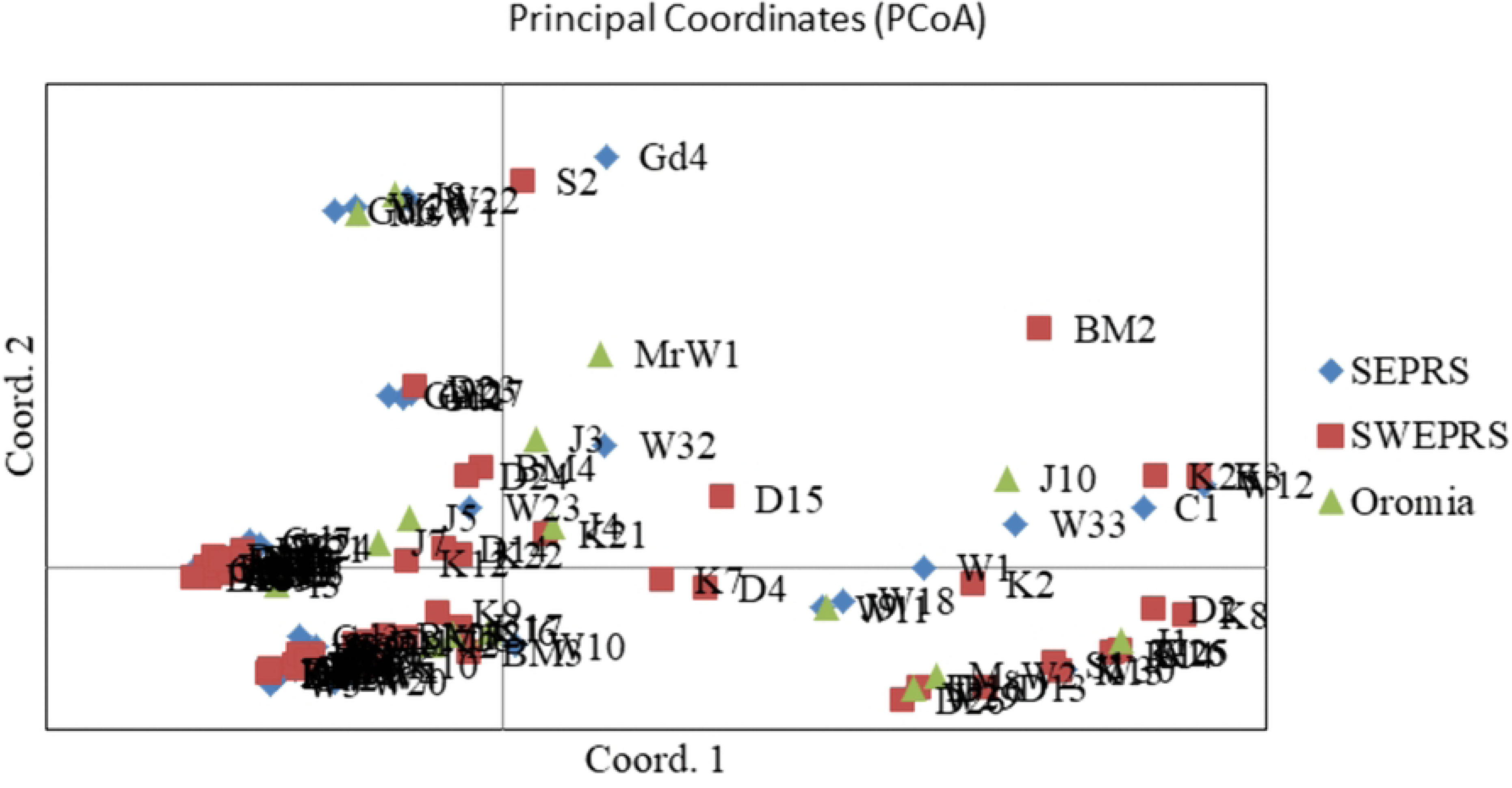
PCoA plot showing genetic relationships among taro accessions based on SSR data. Each point represents an individual, and distances reflect genetic dissimilarity.

### Cluster analysis

NJ and UPGMA analyses were used to examine the relationships among individual accessions. The resulting dendrograms are presented in Figs 5 and 6, respectively.

**Fig 5.**
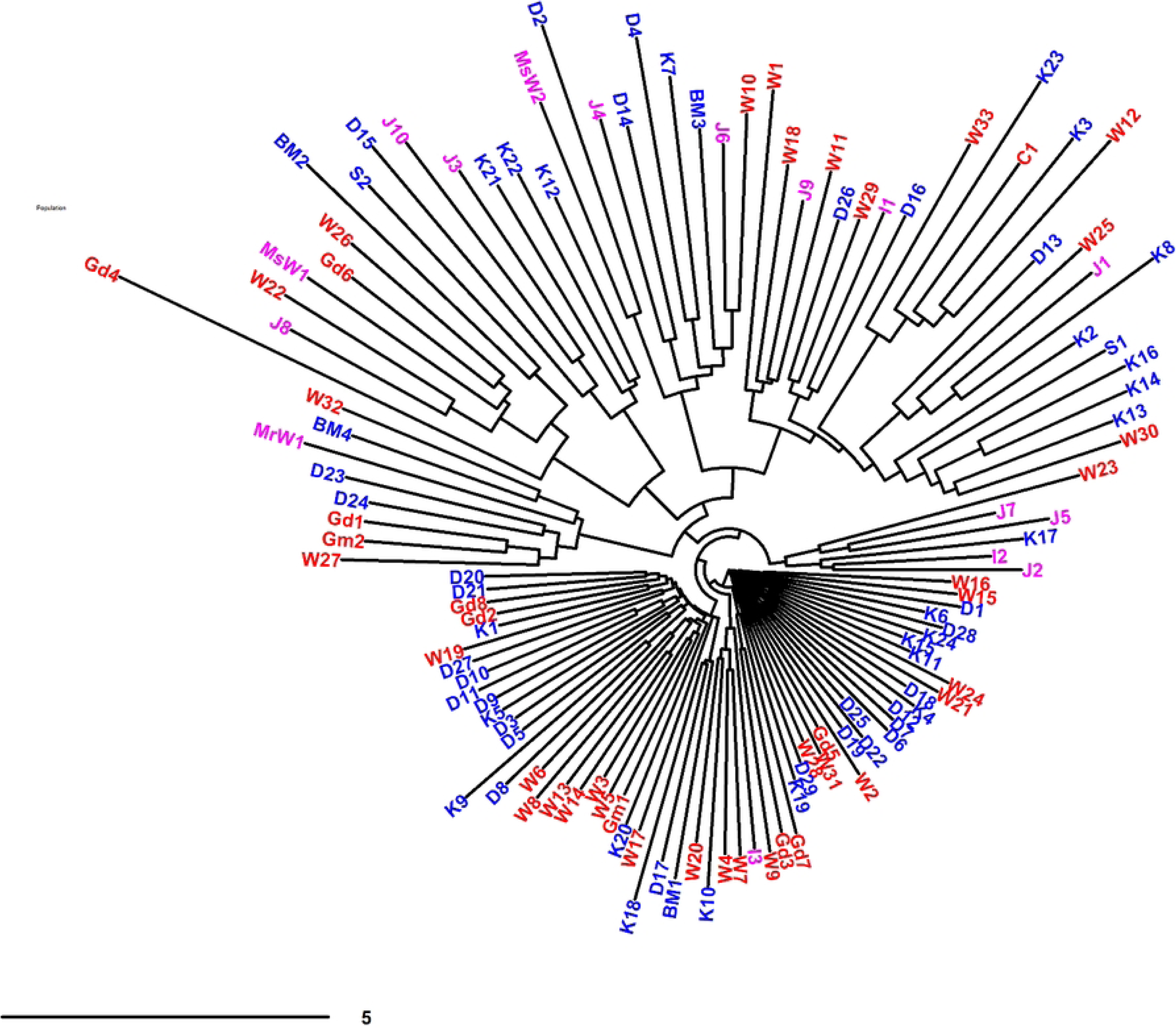
NJ dendrogram of 119 accessions based on genetic distance. Accession labels are colour-coded according to population: SERS (red), SWEPRS (blue), and Oromia (magenta).

**Fig 6.**
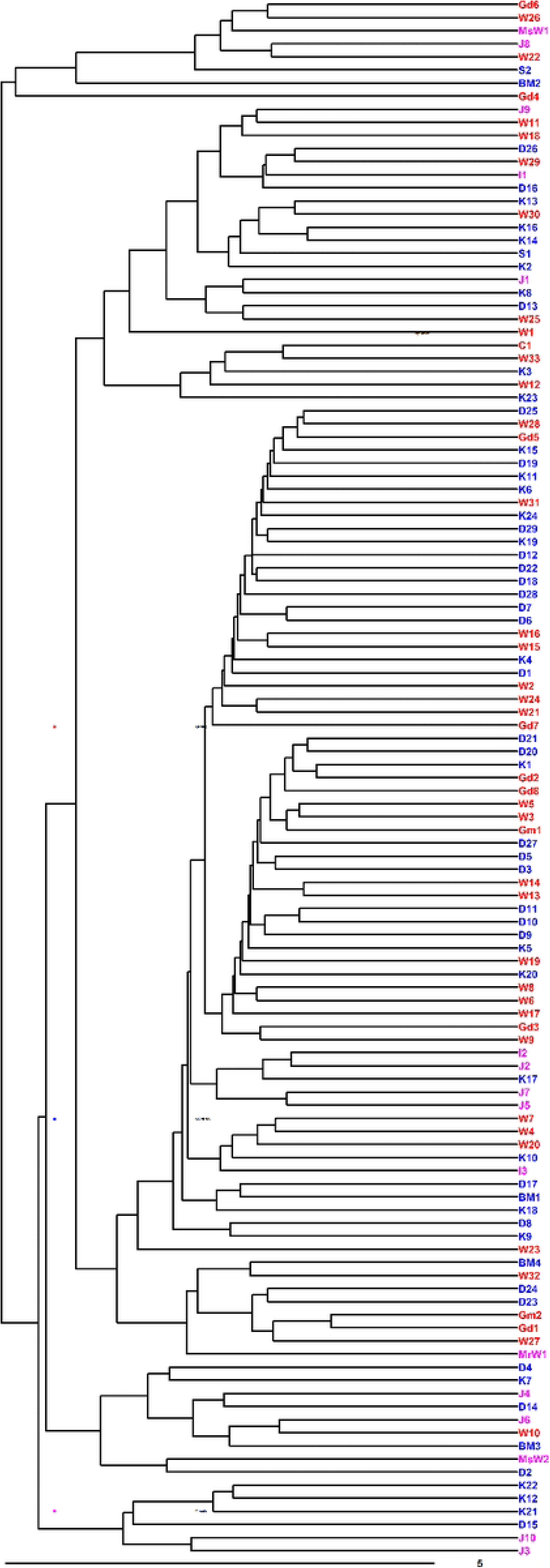
UPGMA phylogenetic dendrogram of 119 accessions based on genetic distance. Accession labels are colour-coded according to regional population: SERS (red), SWEPRS (blue), and Oromia (magenta).

Both clustering methods showed substantial intermixing of accessions from the three regional populations, with no clear clustering according to geographic origin. The clustering patterns were therefore consistent with the extensive overlap observed in the PCoA.

### Population structure analysis

Bayesian population-structure analysis identified K = 2 as the most strongly supported level of genetic structure among the 119 Ethiopian taro accessions. The mean log probability of the data [Ln P(X|K)] increased progressively with increasing K, whereas the Evanno ΔK criterion reached its maximum value of 560.67 at K = 2 (Fig 7A). At K = 2, individual accessions showed varying membership coefficients in the two inferred clusters, with substantial admixture among accessions. The inferred clusters did not correspond clearly to the three geographic populations (Fig 7B).

**Fig 7.**
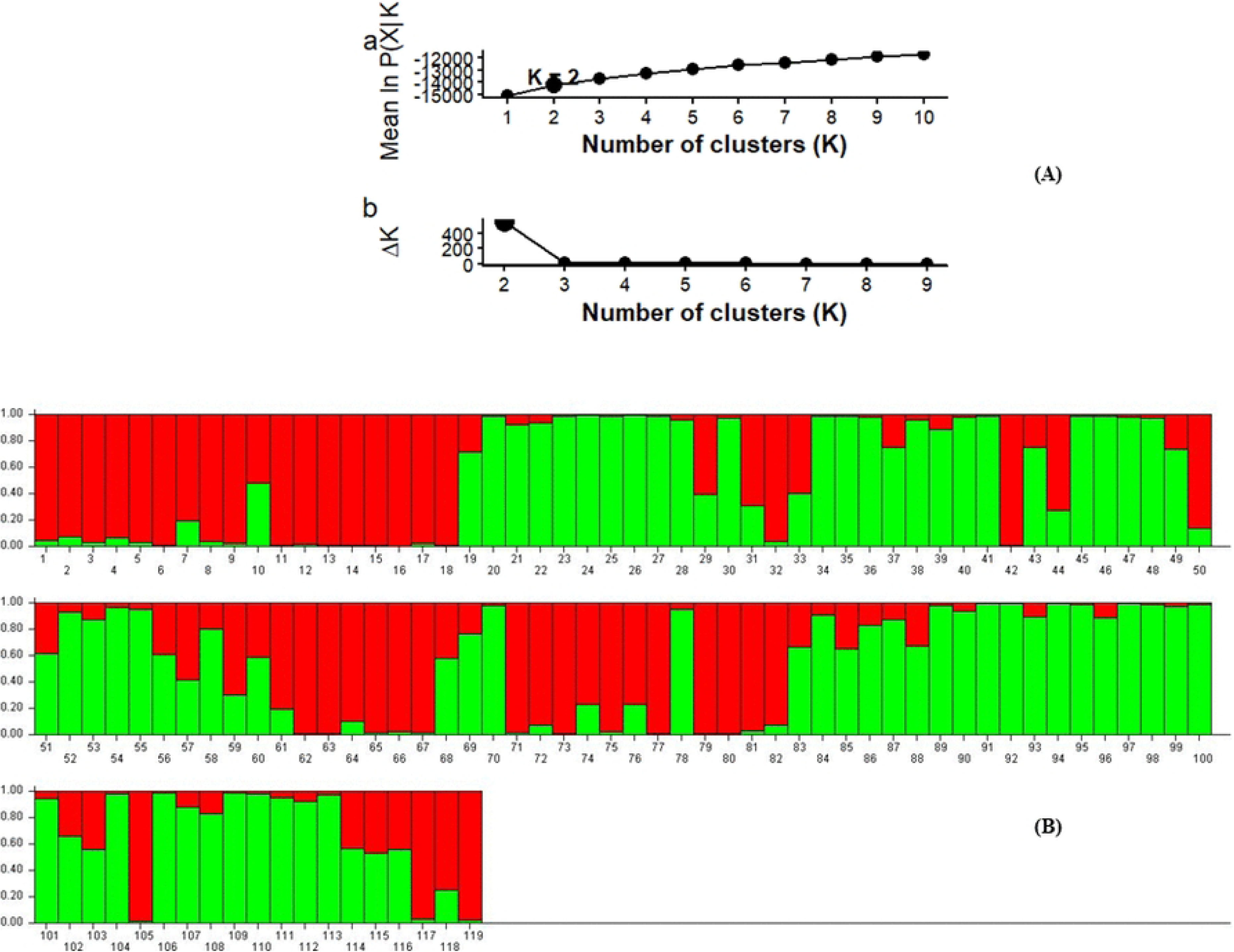
Population structure analysis of 119 Ethiopian taro accessions using STRUCTURE. (A) Evaluation of the most appropriate number of genetic clusters (K): mean log probability of the data [Ln P(X|K)] across five independent runs for K = 1–10 (top) and ΔK values calculated using STRUCTURE Harvester (bottom). The ΔK criterion indicated K = 2 as the most appropriate number of genetic clusters. (B) Population structure inferred at K = 2. Each vertical bar represents one accession, and the y-axis indicates the estimated membership coefficient (Q), ranging from 0 to 1. The two colors represent the inferred genetic clusters, and different color proportions within an accession indicate admixture. Accessions are arranged according to their sampling populations (SERS, SWEPRS, and Oromia). Panel A was generated using R version 4.6.0 from STRUCTURE v2.3.4 output processed with STRUCTURE Harvester.

### Multilocus genotypic diversity and linkage association

Genotypic diversity and multilocus association were assessed using the 199 polymorphic bands scored across the 119 accessions. The population-level estimates are presented in Table 8. All 119 accessions possessed unique multilocus genotypes, with 44, 59, and 16 MLGs identified among the 44 SERS, 59 SWEPRS, and 16 Oromia accessions, respectively. Thus, the number of MLGs equaled the number of sampled accessions in each population.

**Table 8.** Population-level genetic diversity and linkage disequilibrium of 119 Ethiopian taro accessions.

| **Population** | **N** | **MLG** | **eMLG** | **H** | **G** | **λ** | **E.5** | **Hexp** | **Ia** | **r̄d** |
|---|---|---|---|---|---|---|---|---|---|---|
| **SERS** | 44 | 44 | 16 | 3.78 | 44 | 0.977 | 1.000 | 0.1184 | 1.647 | 0.00671 |
| **SWEPRS** | 59 | 59 | 16 | 4.08 | 59 | 0.983 | 1.000 | 0.1091 | 2.176 | 0.00829 |
| **Oromia** | 16 | 16 | 16 | 2.77 | 16 | 0.938 | 1.000 | 0.0911 | 0.902 | 0.00731 |
| **Total** | 119 | 119 | 16 | 4.78 | 119 | 0.992 | 1.000 | 0.1116 | 1.975 | 0.00661 |

*Notes:* N, number of accessions; MLG, number of multilocus genotypes; eMLG, expected number of multilocus genotypes at the standardized sample size; H, Shannon–Wiener index; G, Stoddart and Taylor's index of genotypic diversity; λ, Simpson's index of diversity; E.5, genotypic evenness; Hexp, expected heterozygosity; Ia, index of association; r̄d, standardized index of association. The analysis was based on 119 accessions scored for 199 bands generated by 15 SSR primers.

The Shannon–Wiener diversity index (H) was highest in SWEPRS (4.08), followed by SERS (3.78) and Oromia (2.77). Stoddart and Taylor’s index of genotypic diversity (G) was 59 in SWEPRS, 44 in SERS, and 16 in Oromia. Simpson’s index of diversity (λ) ranged from 0.938 in Oromia to 0.983 in SWEPRS, while genotypic evenness (E.5) was 1.000 in all three populations. Expected heterozygosity (Hexp) ranged from 0.0911 in Oromia to 0.1184 in SERS, with an overall value of 0.1116.

The index of association (Ia) ranged from 0.902 in Oromia to 2.176 in SWEPRS, with an overall value of 1.975. The standardized index of association (r̄d) ranged from 0.00671 in SERS to 0.00829 in SWEPRS, with an overall value of 0.00661.

The overall values of the index of association (Ia) and standardized index of association (r̄d) were 1.975 and 0.00661, respectively. Together with the high multilocus genotypic diversity observed within populations, these results provide additional information on the genetic diversity and multilocus association patterns of Ethiopian taro accessions.

## Discussion

### Marker informativeness and polymorphism

The 15 SSR markers generated 199 reproducible polymorphic loci across the 119 Ethiopian taro accessions, with all scored loci showing polymorphism. The mean PIC was 0.583, with values ranging from 0.300 to 0.888, indicating that the marker set provided substantial discriminatory power for assessing genetic variation in the germplasm analyzed. The relatively high PIC values obtained in the present study contrast with the lower overall PIC reported in the previous analysis and with some earlier studies of taro. For example, [1] reported a mean PIC of 0.43, while [45,46] reported PIC values of 0.65 and 0.428, respectively. Differences among studies may arise from the marker systems used, the number and geographic composition of accessions, the extent of variation within the germplasm, and differences in data scoring and analytical approaches.

The high polymorphism observed in the present study is particularly notable because the SSR profiles were analyzed as dominant presence/absence data owing to the complex ploidy and scoring characteristics of the material. Consequently, the GenAlEx diversity estimates should be interpreted as measures of variation in the scored binary marker states rather than as conventional codominant SSR allele-frequency estimates. Nevertheless, the high PIC values and complete polymorphism across the marker set demonstrate that the selected primers were effective for detecting variation among the Ethiopian taro accessions. The variation in informativeness among primers also indicates that the marker panel captured genetic variation at different levels across the genome.

### Genetic diversity within populations

Despite the complete polymorphism observed across the full collection, the diversity statistics based on the scored binary states were relatively low. Overall expected heterozygosity (He) was 0.066, while population-level He values ranged from 0.047 in Oromia to 0.061 in SERS. Similarly, the effective number of alleles (Ne) showed little variation among populations, ranging from 1.057 to 1.069. These results indicate that, although polymorphic loci were widespread across the collection, the frequencies of the alternative marker states were often uneven, resulting in relatively low effective diversity at individual scored loci.

The population-level results also showed differences in the amount of detectable variation. SWEPRS had the highest mean number of observed allelic states (Na = 1.651) and the highest proportion of polymorphic loci (82.57%), whereas Oromia had the lowest values for Na (0.743), He (0.047), and proportion of polymorphic loci (37.14%). SERS had the highest Shannon’s information index (I = 0.126) and He (0.061). These differences indicate that the three regional populations do not contain identical levels of detectable marker variation, although the magnitude of differentiation among them was small.

The relatively low He values observed in the present study are lower than estimates reported for some Asian taro collections. For example, [34] reported higher diversity in Indian taro using isozyme and random amplified polymorphic DNA (RAPD) markers, while [46] reported an average PIC of 0.428 and expected heterozygosity of 0.485 in Chinese taro based on genome-wide InDel-SSR markers. Such comparisons should, however, be made cautiously because marker systems, genomic coverage, scoring methods, sample sizes, and germplasm composition differ substantially among studies. [4] similarly reported comparatively lower genetic diversity in several African taro collections than in Asian germplasm and discussed the possible contribution of a limited number of introduced clones followed by vegetative propagation and farmer selection.

A similar historical process may have contributed to the relatively low locus-level diversity observed in the Ethiopian material. Taro is predominantly propagated vegetatively, which can preserve farmer-maintained genotypes over long periods and may reduce the contribution of sexual reproduction to the generation of new multilocus variation. However, the present data also demonstrate that low diversity at individual scored loci does not imply a lack of genotypic diversity. All 119 accessions possessed unique multilocus genotypes, demonstrating that different combinations of marker states distinguish the accessions even though variation at individual loci is relatively limited.

The differences in diversity among regional populations should therefore not be interpreted simply as evidence of isolated evolutionary histories. The higher diversity detected in SWEPRS and SERS may reflect the particular composition of the sampled germplasm and the accumulation or maintenance of different planting materials within these regions. Because the overall population differentiation was weak and extensive genetic overlap was observed in the multivariate and clustering analyses, stronger explanations such as long-term geographic isolation or independent population histories are not supported by the present data.

### Population structure, genetic differentiation, and gene flow

The population-level analyses consistently indicated that most genetic variation was distributed within, rather than among, the three predefined regional populations. AMOVA attributed 98% of the total molecular variation to within-population variation and only 2% to among-population variation, with an overall ΦPT of 0.016 (*P* = 0.016). Thus, although the overall differentiation was statistically detectable, its magnitude was weak. This pattern is consistent with previous studies of taro that have reported high within-population variation and relatively weak geographic differentiation, including [31,34,47,48].

The pairwise results provide further support for the weak differentiation observed among the populations. Nei’s genetic distances were extremely small (0.001–0.002), and genetic identities were correspondingly high (0.998–0.999). Significant pairwise differentiation was detected only between SERS and Oromia (ΦPT = 0.038, *P* = 0.018), whereas SERS–SWEPRS and SWEPRS–Oromia were not significantly differentiated. The UPGMA analysis based on population mean genetic distances similarly placed SERS and SWEPRS closer to each other than to Oromia. Importantly, however, this relative difference should not be interpreted as strong population separation because the absolute genetic distances were very small and the overall differentiation among populations was weak.

The estimated Nm values were high for all population pairs, ranging from 6.342 between SERS and Oromia to 24.750 between SERS and SWEPRS. These estimates are consistent with the low ΦPT values and the high genetic similarity among populations. Because Nm is an indirect estimate derived from population differentiation rather than a direct measurement of contemporary movement of propagules, it should be interpreted as evidence of substantial historical or effective connectivity rather than as a direct measurement of current farmer-to-farmer exchange.

The weak geographic structuring was also evident from the PCoA, NJ, and UPGMA analyses. The first three PCoA axes explained only 19.32% of the total variation, and accessions from the three regional populations showed extensive overlap rather than distinct geographic clusters. Similarly, the NJ and individual-accession UPGMA dendrograms showed considerable intermixing of accessions from different regions. Together, these independent analyses indicate that geographic sampling origin does not strongly explain the observed genetic relationships among the Ethiopian taro accessions.

The STRUCTURE analysis identified K = 2 as the strongest partition according to the Evanno ΔK criterion [44] . The NJ tree also appeared to show two major genetic groups, broadly consistent with the STRUCTURE result. However, accessions from different geographic populations were represented in both groups, indicating that the inferred genetic groups were not geographically exclusive. The STRUCTURE barplot further revealed substantial admixture among accessions, and the two inferred groups did not correspond clearly to the three predefined regional populations. Thus, the agreement between STRUCTURE and NJ analyses supports the presence of two broad genetic groupings, while the mixed geographic origins of accessions within these groups indicate weak correspondence between genetic grouping and geographic origin.

The observed pattern is compatible with extensive movement and exchange of vegetatively propagated planting materials among farming communities. Human-mediated movement of taro propagules can introduce genetically similar material into geographically separated production areas and thereby reduce differentiation among populations. Similar weak relationships between genetic structure and geographic origin have been reported in taro by [14,29]. However, the present marker data alone cannot establish the precise historical pathways or current extent of farmer-mediated exchange, and such explanations should therefore be regarded as plausible interpretations rather than demonstrated causes.

### Multilocus genotypic diversity

An important finding of the present study is the contrast between relatively low diversity at individual scored loci and high multilocus genotypic diversity. All 119 accessions possessed unique multilocus genotypes, and the number of MLGs equaled the number of sampled accessions in each regional population. The overall Simpson’s index of diversity was 0.992, while genotypic evenness was 1.000, indicating that the sampled accessions were highly distinct at the multilocus level and that the detected genotypes were represented without substantial repetition.

This pattern demonstrates why the locus-level heterozygosity estimates should not be interpreted as evidence that the Ethiopian taro collection contains little useful genetic variation. Individual loci may show limited variation or unequal frequencies, while combinations of marker states across many loci can distinguish a large number of genotypes. The unique MLGs observed in every accession therefore indicate that substantial genotypic variation is maintained within the sampled germplasm despite the relatively low mean He.

The index of association (Ia) and standardized index of association (r̄d) were 1.975 and 0.00661, respectively, across the complete dataset. These values indicate measurable multilocus association in the dataset; however, without a corresponding permutation test or statistical significance assessment, they should not be interpreted as definitive evidence of linkage disequilibrium. The very high number of unique MLGs nevertheless demonstrates considerable multilocus differentiation among the sampled accessions.

### Implications for conservation and breeding

The results have important implications for the conservation and utilization of Ethiopian taro germplasm. The predominance of within-population variation, together with the very low differentiation among regional populations, indicates that substantial genetic variation is maintained within the sampled populations. Consequently, conservation strategies should give strong attention to intensive sampling within production areas rather than relying exclusively on broad geographic coverage. This is particularly relevant for clonally propagated crops, where individual farmer-maintained accessions may represent distinct multilocus genotypes even when populations show low average differentiation.

The finding that all 119 accessions possessed unique MLGs further emphasizes the value of conserving individual accessions rather than assuming that accessions collected from the same region are genetically redundant. This pattern suggests that substantial genotypic diversity may be maintained through farmer selection, local maintenance of planting materials, and the accumulation of distinct genotype combinations over time. The results are broadly consistent with the importance of within-population variation reported in African taro germplasm by [4].

The close genetic relationship among the regional populations and the high estimated connectivity may facilitate the movement of useful genetic variation among farming communities. At the same time, extensive sharing of planting materials could also result in the broad distribution of genetically similar material and may reduce the geographic distinctiveness of local populations. Therefore, conservation programs should maintain accessions from multiple locations while also preserving individual genotypes that show distinctive multilocus profiles.

For breeding, the high number of unique multilocus genotypes provides a useful starting point for selecting diverse parental materials. The relatively low differentiation among geographic populations suggests that geographic origin alone may not be an efficient criterion for choosing genetically contrasting parents. Instead, multilocus genotype information, combined with agronomic, nutritional, morphological, and other phenotypic traits, could provide a more effective basis for parent selection.

The present findings also demonstrate the value of combining complementary analytical approaches. AMOVA quantified the distribution of molecular variation, Nei’s distances measured relationships among populations, PCoA and tree-based methods visualized relationships among accessions, STRUCTURE identified the strongest statistical partition of genetic variation, and multilocus analysis revealed the high uniqueness of individual accessions. The broadly consistent evidence from these approaches strengthens the conclusion that Ethiopian taro has substantial multilocus genotypic variation but weak geographic population differentiation.

The findings indicate that Ethiopian taro germplasm maintains substantial multilocus genotypic diversity despite relatively low diversity at individual scored loci and weak differentiation among the three regional populations. The predominance of within-population variation and the presence of distinct multilocus genotypes emphasize the importance of conserving individual accessions from both farmer-managed fields and institutional collections. Moreover, geographic origin alone may not reliably identify genetically contrasting materials for breeding, highlighting the value of combining molecular characterization with morphological, agronomic, and nutritional assessments when selecting accessions for conservation and crop improvement.

## Conclusions

This study revealed substantial multilocus genotypic diversity in Ethiopian taro germplasm, despite relatively low diversity at individual scored loci and weak genetic differentiation among the three regional populations. All 119 accessions possessed unique multilocus genotypes, while AMOVA showed that most molecular variation occurred within populations. The STRUCTURE and NJ analyses indicated two broad genetic groupings, but accessions from different geographic populations occurred in both groups, demonstrating that genetic grouping did not correspond closely to geographic origin. These findings emphasize the importance of conserving individual accessions from both farmer-managed fields and institutional collections and of using molecular and phenotypic information together to identify valuable materials for breeding. However, the use of 15 SSR markers scored as dominant binary profiles limits the resolution of allelic diversity and fine-scale genomic relationships. Future studies should employ high-density single-nucleotide polymorphism (SNP) or DArTseq markers and broader, geographically balanced sampling to improve understanding of the genetic diversity, population history, and structure of Ethiopian taro. Integrating these approaches with morphological, agronomic, and nutritional characterization would strengthen the identification and utilization of diverse germplasm for conservation and crop improvement.

## Acknowledgements

We express our sincere gratitude to the Department of Microbial Sciences and Genetics, Addis Ababa University, for laboratory facilities for DNA extraction and quality assessment; Haramaya University for PCR laboratory support; and the Ethiopian Biodiversity Institute (EBI) for providing taro accessions. We deeply appreciate their valuable contributions to this research.

## Supporting information

**S1 Table.** Passport data of Ethiopian taro (*C. esculenta* (L.) Schott) accessions, including local and cultivar names, collection information, geographic coordinates, and altitude.

